# Generating *in vivo* somatic mouse mosaics with locus-specific, stably-integrated transgenic elements

**DOI:** 10.1101/097386

**Authors:** Gi Bum Kim, Marina Dutra-Clarke, Rachelle Levy, Hannah Park, Sara Sabet, Jessica Molina, Aslam Abbasi Akhtar, Serguei Bannykh, Moise Danielpour, Joshua J. Breunig

**Author notes:** Equal contributors (Marina Dutra-Clarke, Rachelle Levy). Correspondence should be addressed to J.J.B.

## Abstract

Viral vectors and electroporation (EP)-mediated gene transfers are efficient means of inducing somatic mosaicism in mice, but they lack the exquisite control over transgene copy number, gene zygosity, and genomic-locus specificity that genetically engineered mouse models (GEMMs) provide. Here, we develop and demonstrate a simple and generalizable *in vivo* method, mosaic analysis by dual recombinase-mediated cassette exchange (MADR). MADR allows for stable labeling of mutant cells express transgenic elements from a precisely-defined chromosomal locus. To test our method, we generated reporter-labeled lineages from stem and progenitor cells in a well-defined *Rosa26^mTmG^* mouse. We demonstrate the power and versatility of MADR by creating novel glioma models with mixed, reporter-defined zygosity or with “personalized” driver mutations from pediatric glioma—each manipulation altering the profile of resulting tumors. Thus, MADR provides a high-throughput genetic platform for the dissection of development and disease, and this rapid method can be applied to the thousands of existing gene-trap mice.

## INTRODUCTION

GEMMs, especially those that enable conditional somatic mosaicism, have been the gold standard for conducting reverse genetics in a temporal-and tissue-specific manner for several decades. Given that GEMM generation is a laborious process, however, many alternative transgenic approaches, such as electroporation (EP)-mediated and viral gene deliveries have been increasingly adapted as a more rapid method of creating somatic mosaics. In EP-mediated gene delivery, for example, a mixture of DNA plasmids can be deposited into the brain ventricular cavity, an electrical current applied, and subsequently the genetic ingredients taken up by the shocked cells. These methods have been used for modeling tumors *in vivo* and lineage-tracing of stem and progenitor cells for developmental studies, illustrating their applicability and versatility (Breunig et al., 2015; Chen and LoTurco, 2012; Friedmann-Morvinski et al., 2012; Hambardzumyan et al., 2011). Moreoever, several researchers have since incorporated into these methods more complex genetic systems, such as tetra-cycline-inducible response elements (TREs), shRNAs, and piggyBac (PB), in order to enable stable and inducible transgenesis and tumor generation *in vivo* (Akhtar et al., 2015; Breunig et al., 2015; Chen and LoTurco, 2012).

Despite offering speed and flexibility, PB-EP and viral methods have some pitfalls. Viral vectors have limited payloads, incite immune responses, and require complex preparation expertise, while both PB-EP and viral delivery suffer from their unpredictable genomic integration patterns, subsequent insertional mutagenesis, and epigenetic transgene silencing (Akhtar et al., 2013; Garrick et al., 1998; Woods et al., 2003). Especially in studies that investigate the role of gain-of-function (GOF) mutations, episomal transfection or transposon-mediated integration often causes significant transgene copy number variability, which can be often inferred by the excessive range of fluorescence brightness, and also chromatin-positional variability, in which some cells can fail to express genes of interest (GOIs) because of unfortunate epigenetic silencing, resulting in clonal genotypic/phenotypic variability (Akhtar et al., 2013; Chandler et al., 2007). In sum, non-GEMM-based evaluation of GOF protein functions is too often confounded by such supraphysiological phenomenon such as overexpression artifacts, unintended cytotoxicity, and transcriptional squelching resulting from copy-number variations (Gibson et al., 2013).

With the unbiased identification of nearly 300 recurrent, putative cancer driver mutations, many of which are GOF oncogenes, it is seemingly imperative to create a tractable *in vivo* platform that can model these potential oncogenes, possibly in conjunction with well-known tumor suppressor LOF tumor genotypes (Lawrence et al., 2014). In contrast to the GOF situation, large-scale knockout (KO) mice consortia now offer facile access to relevant GEMMs for loss-of-function (LOF) mutations in tumor suppressor genes, but even then, creating a multiple-transgenic mouse with several knockout genotypes is significantly time-consuming, and it has been shown that different conditional alleles can recombine independently of each other, potentially mislabeling clones and misrepresenting their genotypes (International Mouse Knockout et al., 2007; Liu et al., 2013). As an alternative, complex CRISPR/Cas9 systems can now simultaneously induce multiple KOs *in vivo* in mice, as well as labeling them (Chen et al., 2015; Zuckermann et al., 2015). For GOF mutations, however, it remains a daunting prospect to compile an exhaustive catalogue of necessary GEMMs.

Given the intrinsic issues with copy number variations, positional variability, and insertional mutagenesis problems of viral and EP-based models, we sought a method that can ensure uniform gene copy number among transfected cells and thus looked to dRMCE, which has been explored as an efficient knock-in method (Lauth, 2002; Osterwalder et al., 2010). Conventionally, this method was explored as a method to modify mouse embryonic stem cells (mESCs) for GEMMs, and often requires antibiotic clonal selection and Southern probing of positive integrants. With appropriate safeguards, we demonstrate that successful dRMCE can be catalyzed *in situ* in somatic cells with tolerable efficiency in a well-characterized reporter mouse *Rosa26^mTmG^ (mTmG)* with definitive genetic labeling of recombined cells (**Fig. 1a**)(Muzumdar et al., 2007). Moreover, we demonstrate the utility of this system in generating mosaicism with a mixture of GOF and LOF mutations, including patient-specific cancer driver mutations. Ultimately, our novel MADR tumor models has a potential to become a first-pass experiment to confirm and study various putative tumor driver mutations and also a rapid pipeline for preclinical drug discovery in a patient-specific manner (Sharpless and Depinho, 2006).

**Figure 1:**
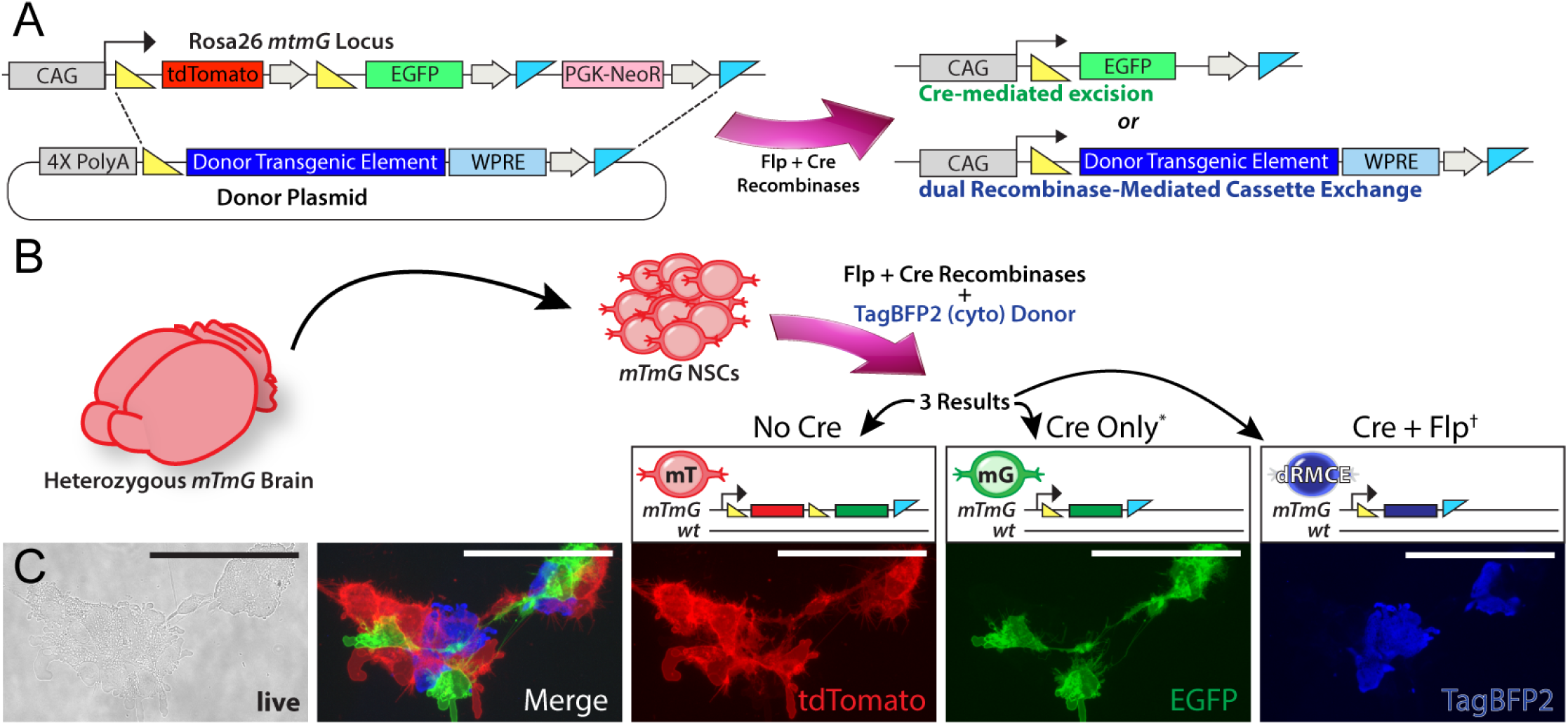
MADR in heterozygous *mTmG* generates three recombinant lineages *in vitro*. A) Flp-Cre expression vector catalyzes either Cre-mediated excision or dRMCE on *Rosa26^mTmG^* allele in the presence a MADR donor vector, resulting in two distinct recombinant products. B) Nucleofection of heterozygous *Rosa26^wT/mTmG^* mNSCs result in three possible lineages: tdTomato+, EGFP+, and TagBFP2+. C) Live imaging of representative cells with non-overlapping fluorescent colors. Scale bars: 100μm

## RESULTS

### Mosaic analysis with dual recombinase donor plasmid strategy and validation

*Rosa26^mTmG^* is a widely used reporter line that constitutively expresses membrane tdTomato and switches to EGFP expression upon Cre-mediated excision of tdTomato cassette (Muzumdar et al., 2007). In order to accommodate MADR-mediated cassette exchange in *mTmG*, we utilized the unused blue fluorescence channel and created a promoter-less donor plasmid encoding *TagBFP2* flanked by loxP and FRT sites (**Fig. 1a**)(Breunig et al., 2015). Notably, both *mTmG* and *TagBFP2* plasmid contain minimal 34-bp FRT, which is refractory to Flp-mediated integration (Lauth et al., 2002), thus circumventing the repeated integration events at the downstream FRT site. Moreover, the open reading frame (ORF) is preceded by PGK polyadenylation signal (pA) and trimerized SV40 pA that will preempt spurious transcription from unintegrated episomes and, more importantly, randomly integrated whole-plasmids, which is known to occur with virtually all means of electrochemical transfections (**Fig. 1a**). The ORF is followed by woodchuck hepatitis virus post-transcriptional regulatory element (WPRE), which increases transgene expression and a rabbit beta-globin pA, which efficiently terminates transcription (**Fig. 1a**). We crossbred Rosa *mTmG* homozygous mice and wild type mice to generate heterozygous *Rosa26^WT/mTmG^* mice (*mTmG^Het^*) and subsequently established a mouse neural stem cell line (mNSC) that carries a single *mTmG* allele. We then induced MADR by nucleofecting the cells with *TagBFP2* donor plasmid (10 ng/μl) and Flp-Cre expression vector (10 ng/μl), which gave rise to 3 distinct genetically labeled lineages, with cells remaining tdTomato+, or turning green or blue (**Fig. 1b,c** and **Supplementary Fig. 1a**)(Anderson et al., 2012). One week after nucleofection with the *TagBFP2* plasmid or a similarly-contructed *TagBFP2-Hras^G12V^* plasmid, FACS analysis gave an approximate efficiency of MADR in mNSCs at around 1% in the case of the *TagBFP2*, and in the case of *Hras^G12V^*, the week-long incubation in the neural stem cell media was long enough to cause integration of the oncogene and the resulting over-proliferation activity (**Supplementary Fig. 1b**). On average, 5% of cells positive for blue fluorescence retained either green or red fluorescence, which can be explained by the relatively slow degradation kinetics of membrane tdTomato, which has been shown to take more than a week to disappear (**Supplementary Fig. 1b,c**)(Muzumdar et al., 2007). After another week of culturing sorted cells, we performed a western blot and confirmed the absence of residual EGFP or tdTomato expression, and also correct *Hras^G12V^* expression, indicating that the recombined *Rosa26* locus generates the correct polypeptide even at the aggregate, polyclonal population level without an antibiotic selection step (**Supplementary Fig. 1d**).

### MADR can be combined with inducible elements to provide orthogonal labeling

Assays for gene function are often performed using transduced or transfected cell lines *in vitro*, but constitutive transgene expression can be detrimental to stable cell line generation, i.e. mutations inhibiting cell cycle or mobility. To obviate this issue, temporally inducible genetic systems, such as TRE, are frequently employed to make the cell line first and then start expressing the GOIs (Akhtar et al., 2015). To expand the utility of single-allele *mTmG^Het^* mNSCs, we aimed to create a pipeline for inducible cell line production by creating a single MADR-compatible plasmid containing rtTA-V10 and TRE-Bi element (**Fig. 2a**)(Akhtar et al., 2015). Subsequently, we generated a colorless TRE-Bi-EGFP cell line with puromycin selection and confirmed the fidelity of TRE with the standard *in vitro* dox treatment (**Fig. 2b,c**).

**Figure 2:**
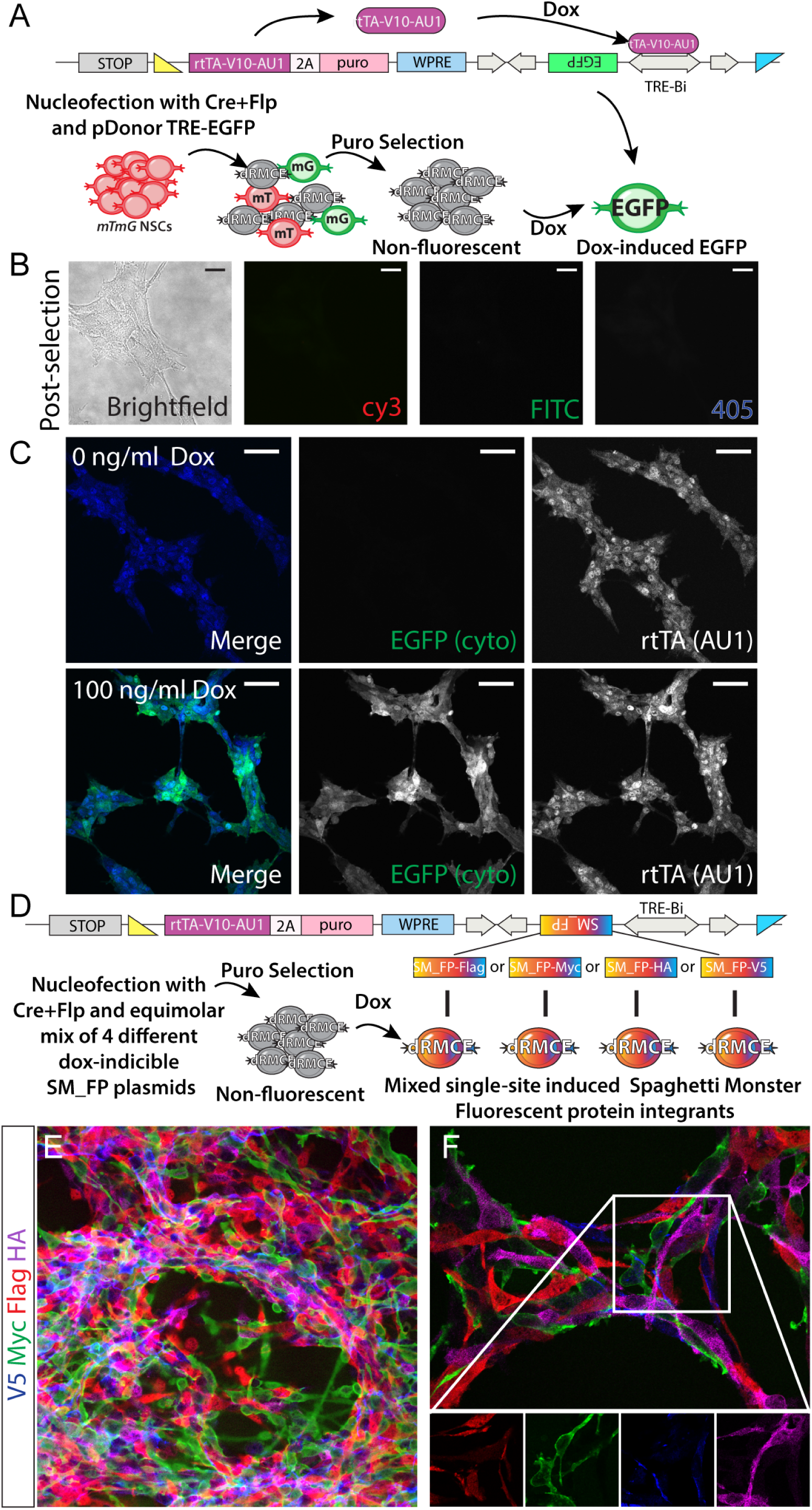
MADR-compatible, selectable, inducible donor construct that can be used for non-leaky reversible expression in vitro and demonstrates single copy integration of pDonor. A) MADR-compatible TRE-EGFP plasmid B) Heterozygous *mTmG* mNSCs are nucleofected with plasmid in a), treated with puromycin, and turned into a colorless population. Scale bars: 10μm C) Induction of EGFP expression in the cell line that constitutively express rtTA-V10-AU1. Scale bars: 50μm D) MADR-compatible TRE-SM_FP plasmids for MADR MAX E) Dox induces efficient SM_FP expression allowing for orthogonal imaging of 4 independent reporters in vitro F) High magnification confocal z section demonstrates that each cell expresses a single SM_FP reporter.

Using a plasmid with puromycin ORF, we attempted to check for the correct recombination at *Rosa26* locus by PCR screening in the cells that expressed puromycin, survived antibiotic selection, and therefore had correct integration at *Rosa26* (**Supplementary Fig. 2a,b** and **Supplementary Table 1**). In those cells that survived puromycin selection after dRMCE step, the tdTomato cassette no longer resides downstream of the CAG-promoter upon dRMCE, which is an important indication that the puromycin-resistant cells were not tdTomato+ cells expressing puromycin from some unknown third chromosomal locations (**Supplementary Fig. 2b, rows 1-2**). However, PCR screening revealed the perdurance of the EGFP cistron in some cells, which is a possible state in very few cells that had Cre-integration but not Flp-excision of EGFP cassette. However, this EGFP cassette is blocked by several polyA elements and situated far downstream from the CAG promoter, which should mitigate any meaningful EGFP expression (**Supplementary Fig. 2b, row 5**). To verify this, we used another plasmid carrying TRE-responsive EGFP element (**Supplementary Fig. 2d**). Using this plasmid and selecting for puromycin-resistant cells, we did not observe EGFP autofluorescence or expression by western blot, and EGFP expression occurred only with doxycycline treatment (**Supplementary Fig. 2c**).

Observing that the antibiotic selection seemed to efficiently kill the EGFP expression in the TRE-Bi-EGFP cell line, we reasoned that we could use similar Dox-inducible plasmids to express four different “spaghetti monster” fluorescent proteins (SM_FPs), which allow for orthogonal detection through their different epitope tags (Viswanathan et al., 2015). We used MADR with multiply-antigenic XFPs (MADR MAX) to empirically assess whether more than one copy of each plasmid could be expressed per cell (**Fig. 2d**). Specifically, expression of more than one of these high signal-to-noise SM_FP probes per cell would be easily detectable by immunofluorescence. Examining hundreds of cells displayed the presence of the SM_FPs in virtually all cells after puro selection and Dox addition, and also in seemingly proportionate ratios (**Fig. 2e**). Furthermore, we did not observe any cells expressing more than one SM_FP by immunofluorescence (**Fig. 2f**), indicating that the dRMCE methodology mediates single copy insertion of transgenic elements.

This *in vitro* system will be beneficial to interrogating GOF protein functions in various primary cell lines derived from any animal carrying loxP and Frt by providing more homogeneous, inducible stable cell lines. As proof-of-principle for this, and to empirically test the utility of the potential leakiness of the 3’ cistron of the TRE-Bi element--which could potentially be activated by upstream promoters or enhancers—we also generated a cell line that inducibly expresses the Notch signaling ligand, Dll1, with a bicistronic TRE-Bi-Dll1/EGFP donor vector (**Supplementary Fig. 2d**). Notably, there was no readily detectable reporter and minute levels of ligand present without Dox; but when added, both EGFP and Dll1 were expressed at virtually similar levels by all cells (**Supplementary Fig. 2d**). (The minute amount of ligand expression in the absence of Dox was comparable to the endogenous expression of mNSCs.) Notch signaling is one example of molecular pathways that are gene-dosage sensitive, and our pipeline could be purposed for studying pathways such as this.

### Efficiency of EP-mediated MADR *in vivo*

To investigate the applicability of *in vivo* MADR, we EP-ed into the neural stem/progenitor cells lining the ventricular/subventricular zone (VZ/SVZ) of postnatal day 2 (P2) *mTmG^Het^* pups with *TagBFP2* donor vector (0.5 μg/μl) and Flp-Cre expression vector (0.5 μg/μl) (**Fig. 3A**). Accordingly, we noted the appearance of recombined cells along the VZ as early as 2 days after EP (**Supplementary Fig. 3A**). By two weeks, many cells differentiated into striatal glia or olfactory bulb neurons (**Fig. 3B, Supplementary Fig. 3B-C**). At this time point, we noticed some rare TagBFP2+ cells with persistant EGFP expression at the VZ, and these cells could be ependymal-lineage cells with slow protein processing kinetics (**Supplementary Fig. 3B**) (Llorens-Bobadilla et al., 2015). However, most TagBFP2+ cells exhibited an absence of tdTomato and EGFP expressions by 2 weeks post-EP (**Supplementary Fig. 3C**). To empirically test the effect of plasmid concentrations on the *in vivo* recombination efficiencies, we varied the concentrations of Flp-Cre recombinase-expression plasmid and MADR MAX reporter plasmid (i.e. expressing a spaghetti-monster reporter plasmid with ten HA-Tags) for high-sensitivity detection of recombined cells (**Fig. 3C** and **Supplementary Fig. 3D**)(Viswanathan et al., 2015). Based on morphological and antigenic comparison of the separate EGFP+ and TagBFP+ populations, it appears that these two different classes of recombinants (EGFP: Excision and BFP: Integration) started out as two separate spatial or temporal populations of precursor cells that favor one reaction over another for unknown reasons. Specifically, we noted that TagBFP+ cells had reduced proportions of striatal glia. Because the MADR (and, thus, MADR MAX) reaction is theoretically irreversible, we examined the brains 2-days post-EP. All DNA mixtures contained a non-MADR, constitutive nuclear TagBfp2 reporter. Surprisingly, we noted that lowering the donor plasmid concentration to 10 ng/μl approached nearly 100% MADR MAX efficiency and almost zero Cre-recombined cells with EGFP expression (**Fig. 3C** and **Supplementary Fig. 3D**). One possible explanation is that increasing the concentration of donor plasmid, hence also of loxP and FRT recombination sites, competes for the recombinases. Alternately, an overproduction of Cre recombinase could exacerbate the Cre-mediated excision and loss of donor plasmids, which means less reactants for the dRMCE reaction. All subsequent electroporation mixtures contained 0.5–1μg/μl of plasmids, in order to generate roughly equivalent numbers of EGFP cells and MADR or MADR MAX cells for side-by-side comparison.

To rule out the possibility that transgene expression was due to the expression from randomly integrated or non-recombined episomes, we performed a series of control electroporations (**Supplementary Fig. 3E**). First, EP of a concentrated mixture of donor *Hras^G12V^* (~5 μg/μl) into wild-type CD1 pups, combined with a PB-EGFP plasmid that marks the lineage of transduced cells, resulted in no abnormal growth, hyperplasia, or tumorigenesis regardless of the presence of Flp and Cre recombinases (**Supplementary Fig. 3E;** for examples of observed phenotypes after MADR of HRas^G12V^ phenotypes see below). In addition, we EP-ed *mTmG* pups with *Hras^G12V^* harboring an inverted loxP and failed to detect any blue recombined cells or hyperplasia by immunostaining, illustrating the specificity of MADR recombination reaction *in vivo* (**Supplementary Fig. 3F**). Several independent EPs of the *Hras^G12V^* donor plasmid and Cre-recombinase alone failed to produce tumor formation when examined at 2 weeks post-EP, indicating that the Flp-excision is an extremely critical step for efficient *in vivo* MADR reaction and Cre is insufficient for sustained transgene integration by itself (Data not shown). (Notably, since Flp-excision is critical to the establishment of irreversible equilibrium, the efficiency of MADR could be further improved by replacing FlpE with FlpO, increasing the efficiency of Flp-exicision (Raymond and Soriano, 2007).) Taken together, these findings suggest that MADR is a highly efficient, locus-specific means of defined copy number transgene manipulation.

**Figure 3:**
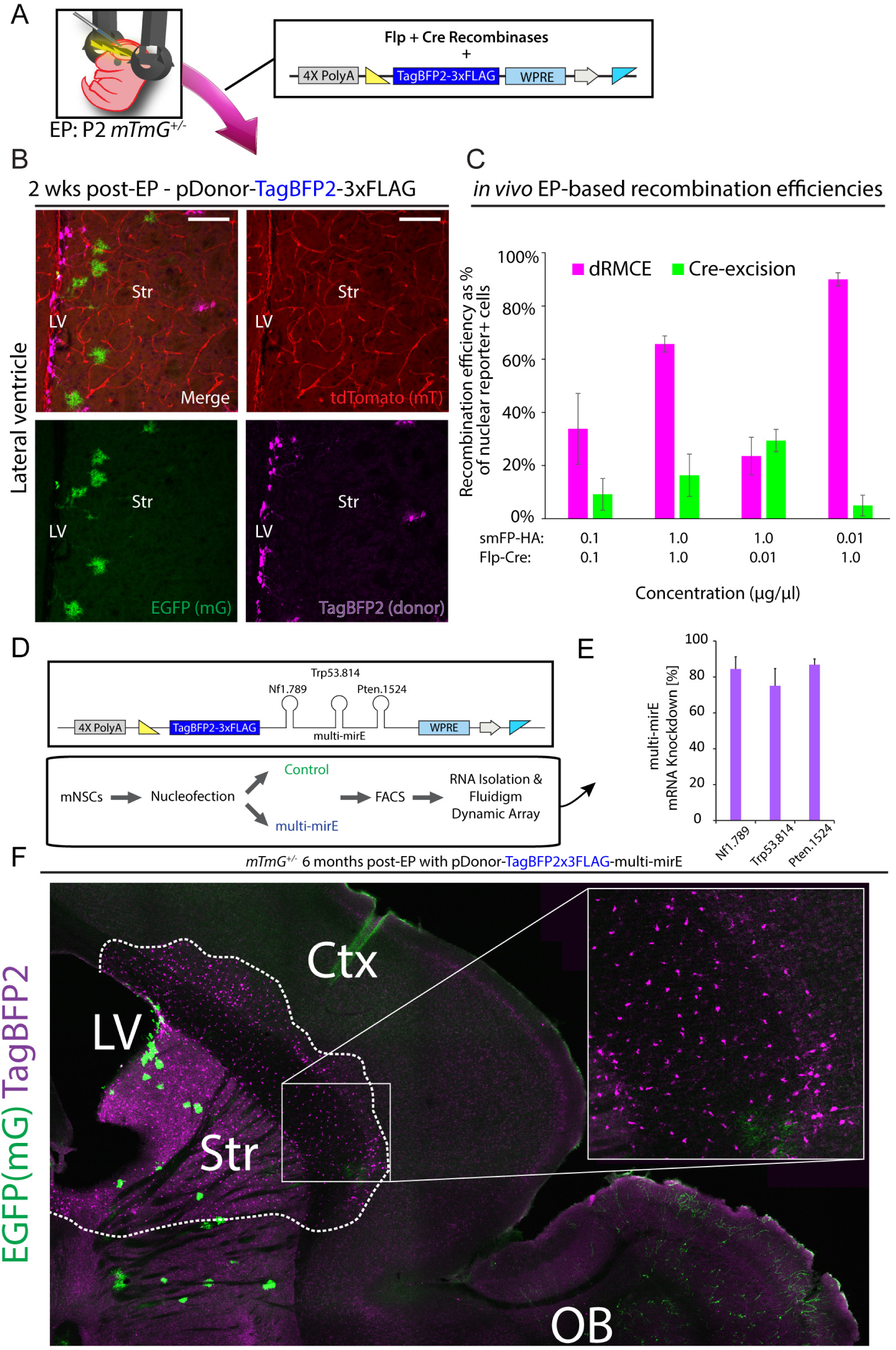
MADR in heterozygous *mTmG* allows for efficient tracing of lineages *in vivo*. A) Standard postnatal electroporation protocol targeting the VZ/SVZ cells in P2 heterozygous *Rosa26^WT/mTmG^* pups with DNA mixture of a Flp-Cre expression vector and a donor plasmid B) Postnatal EP recapitulates *in vitro* nucleofection experiment and yields TagBFP2+ MADR along with EGFP+ and tdTomato+ lineages at 2 weeks post-EP. Scale bars: 100μm C) Electroporation mixtures with different concentration of recombinase and donor plasmids result in various efficiencies of both MADR and Cre-excision recombination reactions *in vivo*. All mixtures contained a nuclear TagBfp2 reporter plasmid. (See **Supp. Fig. 3D** for representative images from this quantitation.) Error bars indicate standard error of the mean (SEM). D) Donor construct for miR-E shRNAs against Nf1, Pten, and Trp53 tied to TagBFP2 reporter E) Validation of knockdown efficacy of multi-miR-E function by qPCR. F) 6-month-old mouse sagittal section showing a hyperplasia of TagBFP2+ cells but no tumor. Scale bars: 200μm and 20μm

Given the ability of our MADR system to independently lineage trace multiple populations of GOF and LOF cells *in vivo,* we reasoned that it might be useful for studying glial development. It was elegantly demonstrated using MADM (Liu et al., 2011) that the combined *Trp53-* and *Nf1-LOF* mutations promote the pre-malignancy hyperproliferation of oligodendrocyte progenitors (OPCs) (Liu et al., 2011). We aimed to mimic a similar developmental phenomenon using our method. First, we created a donor construct harboring three contiguous validated miR-E-based shRNAs targeted at Nf1, Pten, and Trp53 tied to TagBFP2 expression (**Fig. 3D**)(Breunig et al., 2015; Fellmann et al., 2013; Sun et al., 2006). We tested this multi-miR-E construct and observed mRNA-level knockdown efficiency at around 80%, comparable to the originally reported efficiency (**Fig. 3E**)(Fellmann et al., 2013). In agreement with the MADM findings, we observed the selective over-growth of TagBFP2+/Pdgfra+ OPCs, aligning with previous observations that LOF models based on Nf1 and Trp53 result in OPC-driven hyperplasia (**Fig. 3F** and **Supplementary Fig. 3G**)(Liu et al., 2011). Notably, the sibling EGFP+ population, which does not contain the miR-E’s yielded a quantitatively smaller, mixed population of mostly astrocytic cells (**Fig. 3F** and data not shown). These genetically defined (i.e. EGFP+) sibling cells could serve as a useful control cell population in a manner akin to the wild-type cells in MADM GEMM systems (Liu et al., 2011). (MADM and MASTR allow rigorous, *Drosophila*-like deciphering of mutant cells but require de novo GEMM generation depending on the chromosomal location of GOIs (Liu et al., 2011).) We also observed that the expression of three miR-Es did not prevent OB neurogenesis (**Supplementary Fig. 3H**), which is counter to our previous findings using Hras and Errb2 (Breunig et al., 2015), suggesting GOF and LOF mutations resulting in increased Ras/MAPK signaling may lead to subtly different cell fate alterations. Albeit with a small group of animals, we did not detect any malignancy 200 days post-EP, indicating that the complete ablation—rather than knockdown—of any one, two, or all of Nf1 P53 and Pten is likely required for highly penetrant, early-onset tumorigenesis. As confirmation of this, we used CRISPR/Cas9-based knockout of these suppressors. Notably, by EPing a combination of sgRNAs against Nf1, Trp53, and Pten along with SpCas9 and piggyBac-mediated EGFP labeling, we noted the formation of white matter associated, high grade, Olig2^+^ tumors (**Supplementary Fig. 3I-J**) in agreement with GEMM glioma models, the MADM glioma models and an in utero EP-based CRISPR model (Chen et al., 2015; Liu et al., 2011; Persson et al., 2010). Nevertheless, our miR-E studies demonstrate the usefulness of MADR in performing lineage tracing after single-copy, stable knockdown of target genes while providing for an internal “control” lineage (i.e. the EGFP+ Cre-recombined cells).

### Generation of focal glioma models based on *in vivo* MADR

Viral and EP tumor models are increasingly employed to study developmental, evolutionary aspects of cancer in various tissue, but there are issues with various gene delivery methods. We reasoned that our method could be best suited for *in vivo*, autochthonous modeling of putative cancer driver genes, and for proof-of-principle, chose *Hras^G12V^*, a highly-used activating oncogene (Breunig et al., 2015). Histological analysis of growth dynamics in putatively single-copy heterozygous mice indicated that Hras^G12V^ cells rapidly overproliferated when compared with EGFP+ populations (**Supplementary Fig. 4A**). Moreover, we reasoned that we might be able to distinguish between heterozygous and homozygous populations of cells by breeding the *mTmG* mice to homozygosity. Specifically, four possibilities could theoretically result after recombination or insertion and each would have a different combination of genetic markers (**Fig.4A**). After a P2 EP, all of the homozygous *mTmG* EP-ed with *Hras^G12V^* rapidly developed glioma and reached terminal morbidity within 3-4 months (**Fig.4B**; n=4). (Using the same oncogene driven by the CAG-promoter, we have previously shown that PB-EP of *Hras^G12V^* results in 100% penetrant glioma (Breunig et al., 2015).) In homozygous mTmG mice, the MADR reaction was highly efficient even when using 10 ng/μl of plasmid (**Supplementary Fig. 4B**). Interestingly, in our homozygous *mTmG* gliomas, blue-only cells (*Rosa26^HrasG12V×2^*) occupied a bigger patch of tumor cross-section than cells expressing both blue and green (*Rosa26^HrasG12V×1^*) (**Fig. 4C-D**). Previously, *Hras^G12V^* copy number has been shown to confer phenotypic differences, such as growth and apoptosis rates (Beronja et al., 2013). Using PB-EP, we also observe that the brighter EGFP-tagged *Hras^G12V^* cells express phosphorylated Rb1 (pRb1) more than the dimmer EGFP+ cells (**Supplementary Fig. 4C**). Similarly, most of the putatively homozygous *Rosa26^HrasG12V×2^* cells in *mTmG* mice EP-ed with *Hras^G12V^* seem to express pRb1, whereas the hemizygous *Rosa26^HrasG12V×1^* do not (**Fig. 4D-E**). This data points to a possibility that the copy number of oncogenes can significantly alter the profile of resulting tumor populations, as previously observed using GEMMs (Beronja et al., 2013).

**Figure 4:**
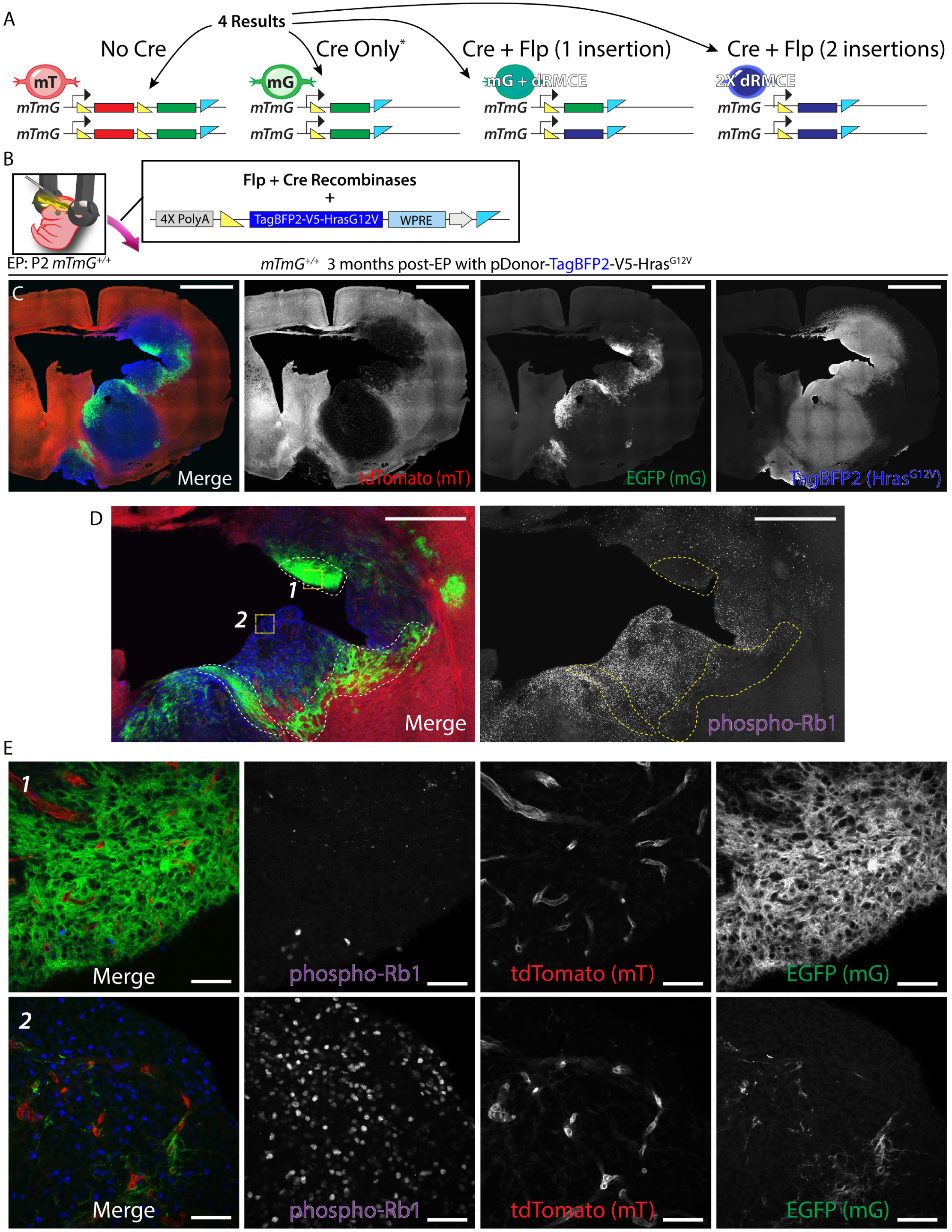
Generation of somatic glioma using *in vivo* MADR with *Hras^G12V^* illustrates that oncogene copy-number can influence the tumor molecular phenotype and growth rate as indicated by end-point tumor volume. A-B) Postnatal EP in homozygous *Rosa26^mTmG^* P2 pups with *Hras^G12V^* oncogene produces two different tumor types (Blue-only *Rosa26^HrasG12V×2^* and blue-and-green *Rosa26^HrasG12V×1^*) C) Representative tumor formation in homozygous *mTmG* 3 months post-EP. Blue-only *Rosa26^HrasG12V×2^* cells occupy a larger section of the tumor than blue-and-green *Rosa26*^*HrasG12Vx1*^. Scale bars: 2mm D) Related to Fig. 2d; Scale bars: 1mm E) Zoom-in images of regions 1 and 2 from b) show phosphorylated-Rb1 expression correlates largely with blue-only cells. Scale bars: 50μm

A current unmet need in terms of mouse models of cancer is a higher throughput method of “personalized” tumor modeling. With the recent surge of knowledge about the putative driver mutations in cancer (Lawrence et al., 2014; Schwartzentruber et al., 2012), there is a need for platforms that can rapidly and comprehensively model these mutations *in vivo* in a combinatorial manner. For example, recently, H3F3A, PDGFRA, and TRP53 were found to be recurrent in pediatric gliomas (Schwartzentruber et al., 2012). Intriguingly, the H3F3A mutations were in two residues—K27 and G34 (Schwartzentruber et al., 2012). Notably, resulting patient tumors bearing either K27M or G34R/V mutations in H3F3A exhibit markedly different transcriptomes (Schwartzentruber et al., 2012) as well as clinical behaviors (Sturm et al., 2012). In particular, patient K27M gliomas cluster along the midline and emerge earlier than G34R/V gliomas, which largely reside in the cerebral hemispheres (Sturm et al., 2012). Nevertheless, H3F3A gliomas of both classes typically harbor TRP53 mutations and can exhibit PDGFRA activating mutations (Schwartzentruber et al., 2012). To model these pediatric tumors in mice, we generated a cassette for co-expression of H3f3a mutations (tagged to EGFP), an activated Pdgfra (D842V), and mutant Trp53 (R270H) (**Fig. 5A**). First, we checked for appropriate expression of H3f3a, Pdgfra, and Trp53 by immunohistochemistry *in vivo* and *in vitro* and noted coincident expression of all proteins (**Supplementary Fig. 5A-B**). Next, we introduced these plasmids by postnatal EP into littermates. Importantly, the electrodes were swept to EP both cortical and striatal VZs to allow for possible tumor formation from both progenitor zones. Fascinatingly, and seemingly in agreement with the clinical presentation of these tumors, K27M-bearing littermates exhibited midline gliomas by P100 (**Fig. 5B**), whereas similarly-treated G34R-bearing littermates mostly displayed diffuse glial hyperplasias and very rare, small tumors (**Fig. 5C**, **arrowhead**). Because G34R mutations often present with ATRX mutations in the second half of the coding sequence (Schwartzentruber et al., 2012), we used CRISPR/CAS9 to introduce InDels mimicking these naturally-occuring mutations. After co-introduction of an SpCas9-expressing plasmid with an sgRNA targeting ATRX along with the G34R-containing co-expression cassette, no gross change was seen in the behavior of G34R glial cells in terms of proclivity for tumor formation at P100 (**Supplementary Fig. 5C**). (A previously-validated antibody recognizing the c terminus of Atrx (Schwartzentruber et al., 2012) demonstrated that >90% of K27M cells and 100% of G34R MADR cells expressed full length Atrx, while >95% of G34R cells with CRISPR/Cas9-targeting of Atrx failed to show Atrx antibody labeling (**Supplementary Fig. 5D-E**).) Notably, at 120 days post-EP, G34R tumors were observed and localized to the cortex and underlying callosum despite equal targeting of the striatal VZ (and coincided hyperplasia of some of these cells; **Supplementary Fig. 5G**). Interestingly, Olig2 signal was significantly more prominent in the dorsal regions, suggesting a cell fate discrepancy between tumor and hyperplasia (**Supplementary Fig. 5G_1_**). At this same 120 day time point, K27M tumors predominantly localized to the sub-cortical structures but cells could be observed in the white matter tracts with a minor amount of cells in the deeper cortical layers (**Supplementary Fig. 5H**). These findings indicate that the K27 and G34 residues are sufficient to significantly regulate the time and location of onset of these glioma subtypes despite the coincident presence of the potent Pdgfra and Trp53 mutations. Further, somatic Atrx mutations do not appear to be a rate-limiting step for tumor formation in G34R tumors.

**Figure 5:**
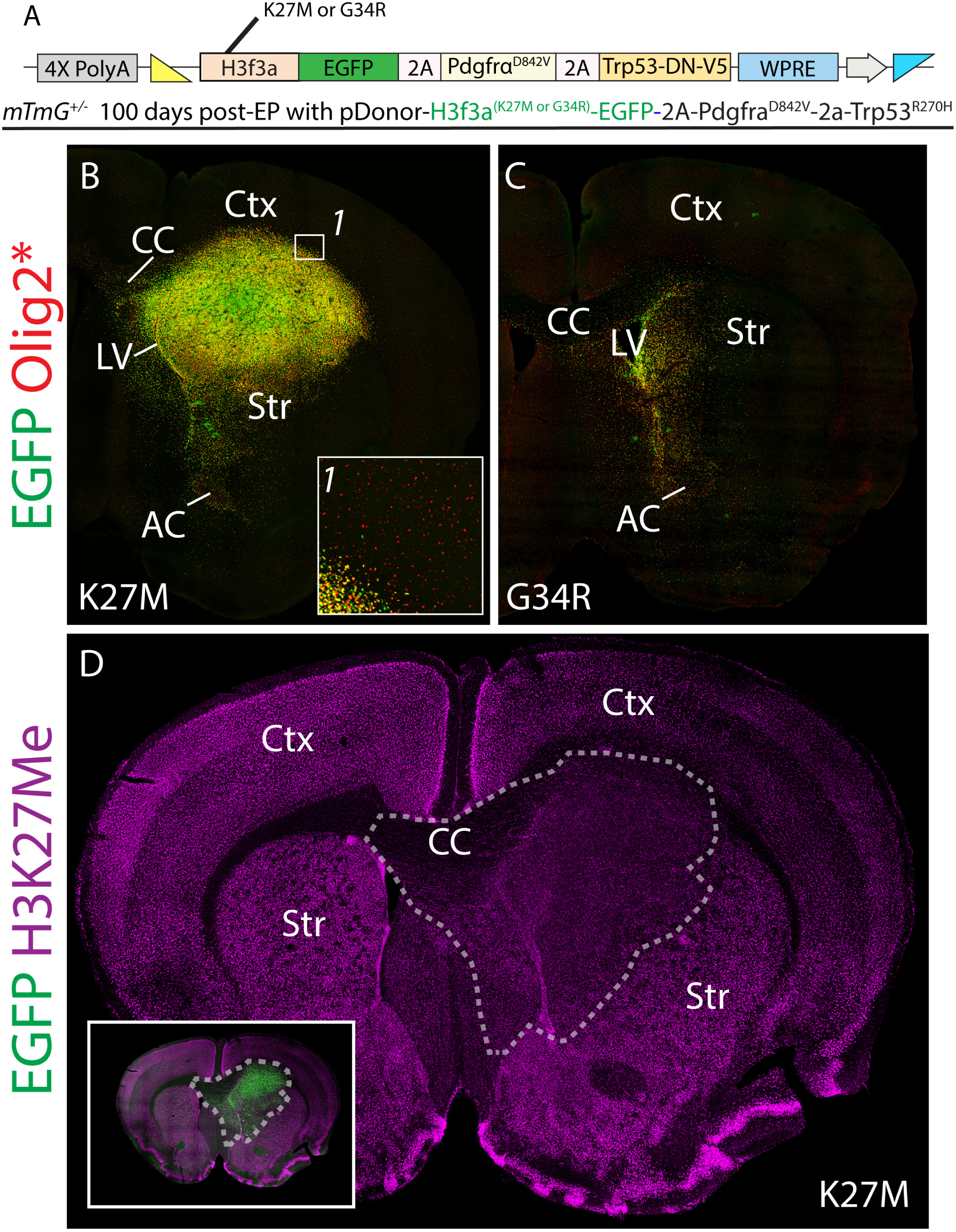
Generation of MADR glioma models utilizing recurrent mutations observed in pediatric GBM yields phenotypes consistent with human subtypes and gives insights into alterations H3f3a PTMs. A) Schematic of donor plasmid for MADR of multiple recurrent pediatric glioma driver mutations B) Representative tumor formation in heteroygous *mTmG* 100 days post-EP. Nuclear EGFP+ *Rosa26^H3f3a-K27M/Pdgfra/Trps3^* cells form a large striatal tumor. Inset B-1 shows a lack of significant cortical infiltration. C) A littermate *Rosa26^H3f3aG34R/Pdgfra/Trp53^* exhibits a glial hyperplasia in the striatum and a small mass of EGFP+ cells in the ventral forbrain medial to the piriform cortex. D) *Rosa26^H3f3aG34R/Pdgfra/Trp53^* EGFP+ tumor cells are hypomethylated at H3K27.

Mechanistically, it is thought that K27M mutations lead to hypomethylation at this residue. In fact, given the intrinsic ability of MADR to lineage trace tumor cells, we were able to confirm the hypomethylation of K27M mutant cells by K27Me antibody (**Fig. 5D**). (This was not simply an artifact of tumor growth as CRISPR/Cas9-mediated knockout of Nf1/Trp53 led to the formation of glioma that was hypermethylated (data not shown).) Taken together, the ability to easily and unambiguously observe such post-translation changes in a tumor cell autonomous manner in vivo holds great promise for the future investigations of disease pathomechanisms in these and other cancers.

### Lineage tracing CRISPR/Cas9 induced MADR MAX glioma models

Recently, CRISPR/Cas9 has been demonstrated to be highly efficacious for the mutation of genes in vivo using EP (Chen et al., 2015; Zuckermann et al., 2015). However, a shortcoming of these studies is a definitive way to lineage trace modified cells. To address this issue, we created a SM_BFP2-P2a-SpCas9-containing donor plasmid to simultaneously label and mutate cells, enabling faithful tracing of mutant cells *in vivo* (**Fig. 6A**). Given our aforementioned findings with the tandem miR-E’s and the Cas9-mediated knockout of Nf1, Trp53, and Pten, we used these same PCR-ed sgRNAs to target Nf1 and Trp53. Successful targeting in EPed cells was confirmed in tumor population gDNA by sequencing (**Fig. 6B**). At roughly 5 months we observed terminal morbidity in EPed animals. Pathological analysis led to the diagnosis of glioblastoma multiforme, principally due to the presence of necrosis in the tumor. Immunohistochemically, we observed that the tumor was largely devoid of TdTomato-labeled populations with the exception of vasculature (**Fig. 6C**-**C**_**1**_, **Supplementary Fig. 6A**). A small EGFP population was observed near where the original targeting site was expected to reside (**Fig. 6 C**, **C_2_**; arrowhead). However, the tumor was demarcated by MADR MAX SM_BFP-V5 labeled cells, which had overgrown the hemisphere (**Fig. 6C**, **C_3_**). Most of this volume was filled with Olig2+ populations though regions lacking signal were observed (**Fig. 6D**; arrowhead; **6E**). As Olig2 is a marker of proneural glioma subtypes, which we and others have observed to precede mesenchymal evolution, we assessed whether this might be a site of tumor evolution by staining for the mesenchymal marker, CD44. Notably, CD44 was found throughout the tumor but was enriched in this Olig2-diminished region (**Fig. 6E**-**F** and data not shown). These results demonstrate the ability of MADR MAX to lineage trace CRISPR/Cas9-mediated knockout of tumor suppressors, leading to glioma formation.

**Figure 6:**
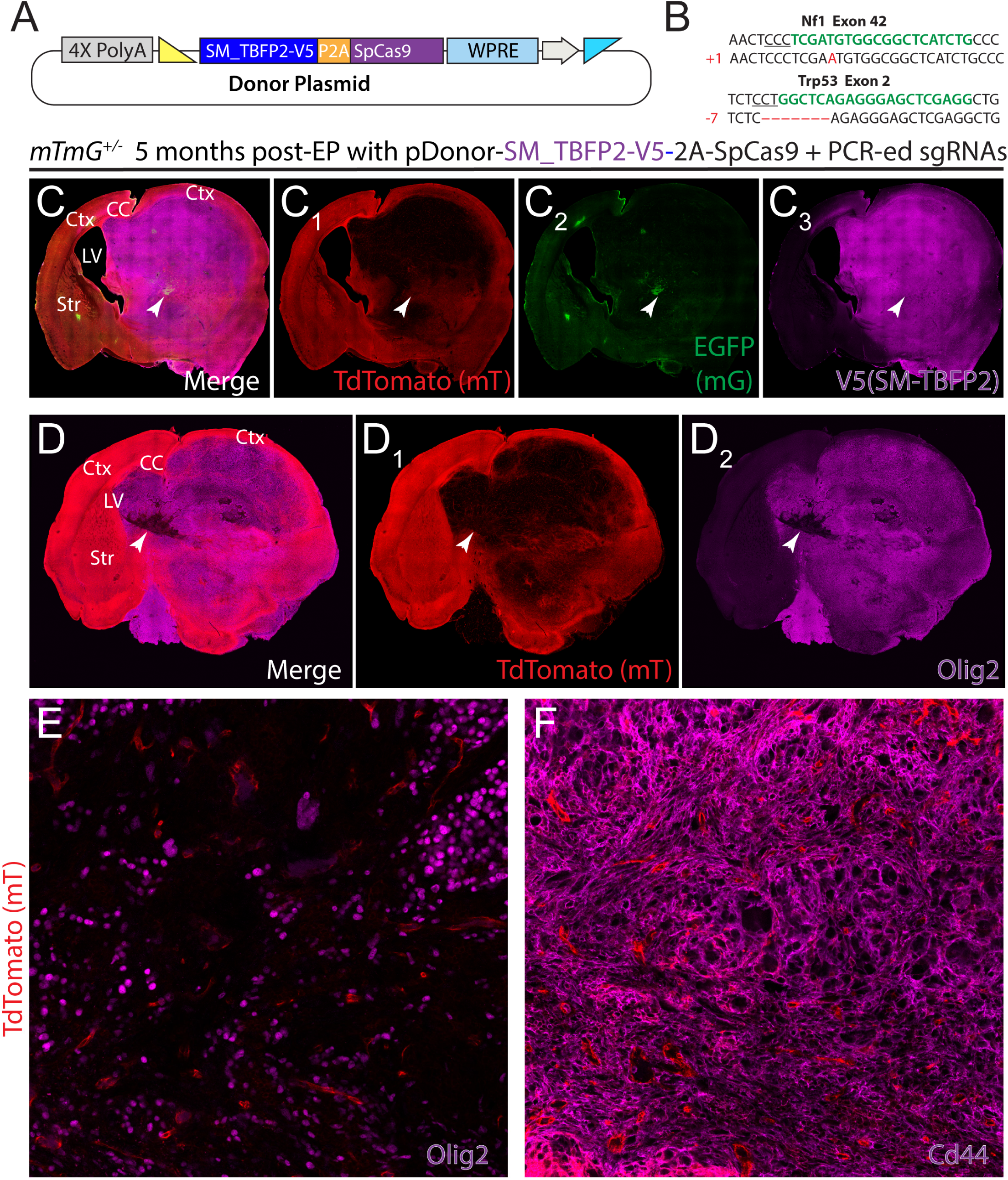
Rapid generation of somatic mosaics using *in vivo* MADR-incorporated Cas9 and PCR-derived sgRNAs allows for interrogation of transdifferentiation within glioma. A) Plasmid for MADR of a TagBFP2-V5 reporter protein and SpCas9 B) tdTomato-/EGFP- glioma cells purified from tumor exhibit InDels in Nf1 and Trp53. C) MADR insertion of TagBFP2-V5 reporter and Cas9 with co-EPed PCR-derived sgRNAs yields high grade glioma observable through genetic labeling of 3 recominant lineages D) Glioma cells are largely Olig2+ with small pockets of heterogeneity (white arrow). E) High magnification Olig2 and tdTomato image focusing on the region denoted by the white arrow in **Fig. 6D**. F) CD44 and tdTomato immunostaining in a roughly adjacent section and region from **Fig. 6E** demonstrating positivity for the CD44 mesenchymal tumor marker.

## DISCUSSION

We demonstrated that MADR-mediated somatic transgenesis *in vivo* is a highly robust method for creating numerous somatic mosaics with a manageable amount of crossbreeding. One drawback using our method is that multiple Cre-mediated integration reactions can occur on wild-type loxP as long as Cre expression lasts, but Cre-mediated stable integration is not favored, as demonstrated by the lack of *Hras+* tumor when the donor *Hras* plasmid was electroporated with Cre recombinase alone, and the episomal plasmids are lost very quickly (Akhtar et al., 2015). Additionally, our safeguards ensure only one definite CAG-promoter-driven in-frame ORF. Our *in vivo* MADR efficiency experiment indicates that diluting the concentration of donor construct possibly mimics single-substrate-based recombination reactions that occur in double-transgenic GEMMs with floxed loci and Cre-recombinase (**Fig. 3C**). We demonstrated the potential to combine two broad modes (GOF/LOF) of tumor mutations expeditiously and flexibly. All in all, we have introduced a new technique to produce stable somatic mosaicism in mice with EP. The intrinsic features of MADR compared favorably with existing methods for in vivo mouse genetic manipulation using other methods (**Table 1**). There is a continued interest in *in vivo* generation of somatic mosaics, as demonstrated by the recent study showing the possibility of transfecting post-mitotic cells *in situ* with the nuclear pore dialator trans-cyclohexane-1,2-diol to the plasmid mixture (De la Rossa et al., 2013). Additionally, viral vectors could be adapted to transduce MADR constituents.

**Table 1:** Comparison of approaches for genetic manipulation in the brain

|  |  |  |  |  |  |
| --- | --- | --- | --- | --- | --- |
| Method | GEMM | Standard EP | Transposition-mediated EP | Virus | MADR |
| Time for engineering and generation | Months | ~2 weeks per plasmid | ~2 weeks per plasmid | >4-6 weeks | ~2 weeks per plasmid |
| Copy number | 1-2 per knock-in | Highly Variable | Highly Variable | Variable but likely less than EP | 1-2 depending on zygosity of recipient |
| Breeding | More complex for conditional alleles | Not necessary | Not necessary | Only necessary for RCAS/Tva | 1 line per targeted strain |
| Stability of Expression | Generally stable depending on locus silencing | Prone to dilution and/or silencing | Prone to silencing and insertional effects | Prone to silencing and insertional effects | Generally stable depending on locus silencing |
| Payload | Limited by targeting construct* | Typically governed by plasmid limits* | Typically governed by plasmid limits* | Limited to viral payloads | Typically governed by plasmid limits* |
| Focality | Depends on cis regulatory elements | Focality depends on electrode orientation | Focality depends on electrode orientation | Diffusion pattern unidirectional from injection site | Focality depends on electrode orientation |

\*-BAC DNA can be utilized
See text for further details

### New MADR EP toolkit

Transposons, such as PB and Sleeping Beauty, are increasingly used in combination with EP to produce stable somatic trangenics and allow long-term developmental studies and also *in vivo* tumor generation (Breunig et al., 2015; Chen and LoTurco, 2012; Chen et al., 2015; Glasgow et al., 2014; LoTurco et al., 2009). Our new method overcomes transposon-based systems’ two major problems: random genomic insertions and copy number variability. Several cell-fate determinants have been shown to result in dramatically differential phenotypes based on their expression levels. High Nfia expression in glial progenitors favor their differentiation into astrocytes, while low expression is observed in cells that become oligodendrocytes, and in another example, higher Fezf2 expression induces the NSC quiescence in the VZ/SVZ by upregulating Notch signaling effectors (Berberoglu et al., 2014; Deneen et al., 2006). Notch signaling is one key example of gene-dosage-sensitive molecular pathways (Ables et al., 2011). Using one-copy vs. two-copy comparison paradigm shown in this report, we demonstrate the potential to study such genetic determinants in one mouse with the ability to compare one-copy, two-copy, and intra-section control cells.

### Extensibility of MADR to existing GEMMs

This strategy can be employed with any off-the-shelf GEMM harboring dual recombinase sites, specifically the IKMC mice with thousands of transgenic mice that already harbor loxP and FRT sites around loci of interest (e.g. Ai14, R26-CAG-LF-mTFP1, Ribotag, and IKMC mice) (**Supplementary Fig. 7**) (Imayoshi et al., 2012; International Mouse Knockout et al., 2007; Madisen et al., 2010; Sanz et al., 2009). Like *Rosa26^mTmG^*, Ai14 is another widely employed reporter mouse line that are typically crossbred with recombinase-expressing transgenic mice (Madisen et al., 2010). By appropriately orienting the recombinase recognition sites, as indicated in **Supplementary Fig. 7**, donor plasmids can be created for use in Ai14 mice, as well as other compatible mice. Ribotag use Cre recombination to swap an untagged Rpl22 Exon 4 with an HA-tagged variant for affinity immunoprecipitation of Cre expression-defined mRNA (Sanz et al., 2009). With MADR, an alternate tag can be inserted with additional elements to allow for simultaneous orthogonal mRNA purification using sequential immunoprecipitation for the tags. In addition, one can create an ORF that begins with a splice acceptor and use this construct to investigate the effects of substituting transgenes under foreign cis-regulatory environments (**Supplementary Fig. 7**). At the same time, a donor plasmid with a fluorescent reporter can be simply electroporated (with Flp and Cre) to enable lineage-tracing from a focal point without the need for Cre-expressing mice in various transgenic mice that flank important loci with loxP and FRT sites.

Because our method requires two different recombinases, one can also drive the expression of these recombinases with different combinations of promoters to restrict the types of cells that are recombined. Finally, there is the exciting possibility of *in vivo* MADR with large-cargo bacterial artificial chromosomes (BAC). A donor plasmid harboring large chunks of genomic fragments driving the expression of fluorescent reporter or recombinases, such as vCre or sCRE, can be created with loxP and FRT sites added on each end, enabling further higher-order lineage tracing studies (Fenno et al., 2014). Next generation sequencing has exponentially increased our understanding of the genomic and transcriptomic changes that occur in tumorigenesis. However, with the catalogue of recurrent somatic mutations in tumors continuing to grow, an emerging problem is to separate tumor-promoting driver mutations from passenger mutations. Further, it is now increasingly appreciated that similarly histologically classified tumors can often have disparate genetic underpinnings that create notably different tumor phenotypes (e.g. K27M vs. G34R tumors). GEMMS have been critical for the understanding of glioma development (Liu et al., 2011; Persson et al., 2010). In particular, the elegant MADM methodology has provided unique insights into the tumor cell of origin and potential treatments (Persson et al., 2010). However, creating the appropriate MADM mice for modeling all of the recurrent mutations and their subsequent combinations would be a daunting task. We show proof of principle for using MADR as a platform for rapid ‘personalized’ modeling of pediatric GBM. By combining MADR GOF transgenesis, and CRISPR/Cas9 LOF manipulations, it is possible for a small lab to generate the plasmid reagents necessary to cover the spectrum—and, thus, possible combinations—of mutations for most tumor types. Recently, CRISPR/Cas9 has been introduced by electroporation along with donor oligonucleotides and plasmids to allow for homology-dependent repair (HDR)-mediated insertion of transgenic elements (Mikuni et al., 2016). This strategy has the advantage of allowing for genome-wide targeting of any transgenic element. However, the percentage of appropriate insertions was many fold below what MADR allows and as with most HDR strategies, insertion will likely be inversely correlated with donor size.

Our findings therefore establish in vivo MADR as a robust methodology for stable mosaic analysis, one which overcomes many of the inherent drawbacks in viral, GEMM, EP, and transposon-based approaches. Additionally, this genetic framework is adaptable to the thousands of strains of mice engineered with dual recombinase recognition sites. Thus, these tools promise to allow for efficient, higher throughput investigation of gene function in development and disease.

## EXPERIMENTAL PROCEDURES

All mice were used in accordance with the Cedars-Sinai Institutional Animal Care and Use Committee.

### Plasmid Cloning

Flp-Cre recombinase expression constructs were generously provided by Y. Voziyanov and previously validated in the context of dRMCE (Anderson et al., 2012). Our donor plasmids were derived from PGKneotpAlox2, using In-Fusion cloning (Clontech) in combination with standard restriction digestion techniques (Soriano, 1999). Briefly, FRT site was created by annealing two oligos and infusing the insert into PGKneotpAlox2. Downstream generation of donor plasmids were done by removing the existing ORF and adding a new cassette using In-Fusion or ligation, as was done for the smFP-HA ORF (Addgene 59759). PB-CAG-plasmids were previously described and created using combination of In-Fusion and ligation strategies (Breunig et al., 2015). Primer sequences used for In-Fusion reactions are available upon request. PCR was done using a standard protocol with KAPA HiFi PCR reagents.

### Mice and electroporation

*Gt(ROSA)26Sor^tm4(ACTB-tdTomato-EGFP)Luo^*/*J* mice (JAX Mice) and were bred with wild-type CD1 mice (Charles River) to generate heterozygous mice. Postnatal lateral ventricle EPs were performed as previously described (Breunig et al., 2015). P1-3 pups were placed on ice for ~5 min. All DNA mixtures contained 0.5-μ1g/μl of Flp-Cre expression vector, donor plasmid, hypBase, or CAG-reporter plasmids diluted in Tris-EDTA buffer, unless noted otherwise. Fast green dye was added (10%v/v) to the mixture, which was injected into the lateral ventricle. Platinum Tweezertrodes delivered 5 pulses of 120 V (50ms; separated by 950 ms) from the ECM 830 System (Harvard Apparatus). SignaGel was applied to increase conductance. Mice were warmed under a heat lamp and returned to their cages.

### Tissue preparation

After anesthesia, mouse brains were isolated and fixed in 4% PFA on a rotator/shaker overnight at 4°C. Brains were embedded in 4% low-melting point agarose (Thermo Fisher) and sectioned at 70 μm on a vibratome (Leica).

### Immunohistochemistry

Immunohistochemistry was performed using standard methodology as previously described (Breunig et al., 2015). Agarose sections were stored in PBS with 0.05% sodium azide until use. Details on the primary antibodies can be found in **Supplementary Table 2**. All primary antibodies were used in PBS-0.03%Triton with 5% normal donkey serum. All secondary antibodies (Jackson ImmunoResearch) were used at 1:1000.

### Cell culture and nucleofection

Three heterozygous P0 *mTmG* pup brains were dissociated to establish the mouse neural stem cell line used in the study. The cell line was maintained as previously described (Breunig et al., 2015). Cells were grown in media containing Neurobasal®-A Medium (Life Technologies 10888-022) supplemented with B-27 without vitamin A (Life Technologies 125 87–010), GlutaMAX (Life Technologies 35050), Antibiotic-Antimycotic (Life Technologies 15240), hEGF (Sigma E9644), heparin (Sigma H3393), and bFGF (Millipore GF003). Neural stem cell nucleofection was performed using the Nucleofector 2b device and Mouse Neural Stem Cell Kit according to manufacturer’s recommendations (Lonza AG). The nucleofection mixtures contained plasmids with equal concentrations of 10 ng/μl.

### Imaging and processing

All fixed images were collected on a Nikon A1R inverted laser confocal microscope. The live image of mNSCs was obtained on an EVOS digital fluorescence inverted microscope. For whole brain images, the automated stitching function of Nikon Elements was used. ND2 files were then imported into ImageJ to create Z-projection images, which were subsequently edited in Adobe Photoshop CS6. In **Fig. 3F**, rotation led to colorless space in the area outside of the tissue section and black fill was added. Adobe Illustrator CS6 was used for the final figure production.

### Quantification of *in vivo* MADR efficiency

For each condition, two pups were electroporated with pCAG-TagBFP2-nls, pDonor-smFP-HA, and Flp-2A-Cre. The brains were taken two days post-EP, and two non-adjacent sections from each brain were stained with HA-Tag antibody and EGFP. For each section, ~25 BFP+ cells were randomly selected, among which HA+ and EGFP+ cells among BFP+ cells were counted. The proportions were averaged over four sections for each group.

### Flow cytometry

Cells were collected as previously described (Breunig et al., 2015). Cells were briefly rinsed in PBS, removed by enzymatic dissociation suing Accutase (Millipore), pelleted at 250g for 3 min, and resuspended in the media. FACS was done on a Beckman Coulter MoFlo at the Cedars-Sinai Flow Cytometry Core.

### Western blot

The cell pellets were resuspended in laemmli buffer and boiled for 5min at 95°C. Protein concentrations were measured on a ThermoScientific NanoDrop 2000. After SDS-PAGE separation and transfer onto nitrocellulose membranes, proteins were detected using the antibodies listed on **Supplementary Table 2**, diluted in 5% milk in 0.1% PBS-Tween. All secondary antibodies (Li-cor IRDye®) were used at 1:15000. Infrared detection was accomplished by the Li-Cor Odyssey^®^ CLX Imaging System.

### Doxycycline and puromycin administration

Doxycycline (Clontech 631311) was added to culture media at the final concentration of 100ng/ml. Puromycin (Clontech 631305) was used at 1μg/ml.

### Multi-miR-E knockdown efficiency quantification

We have previously used FlEx-based transgene expression, specifically Cre-mediated inversion and activation of EGFP cassette (FlEx-EGFP)(Breunig et al., 2015). To test our multi-miR-E targeting Nf1, Pten, and Trp53, we made a CAG-driven FlEx-based construct harboring the multiple miR-Es (FlEx-multi-miR-E). Postnatal mNSC line was established by dissociating CD1 pup brains, transfected with EGFP or FlEx-multi-miR-E and Cre-recombinase vector. Fluorescent cells were sorted and subjected to mRNA extraction and SYBR-based Fluidigm BioMark dynamic array using qPCR probes for Nf1, Pten, and Trp53.

### PCR-generation of U6-sgRNA fragments

A short reverse primer and an ultramer forward primer (IDT DNA) were combined in a PCR reaction and subsequent purification to make concentrated sgRNAs (Ran et al., 2013). 100ng of each fragment was combined with plasmid DNA for EP.

### Sequencing InDel mutations in murine tumor cells

A pure population of tumor cells was obtained by FACS and genomic DNA was isolated (Qiagen DNeasy). Using primers flanking the gRNA target site, we PCR amplified the regions expected to contain InDel mutations for NF1, Trp53, and Pten. The PCR amplified fragments were topo cloned using the Thermo Fisher Zero Blunt TOPO kit and transformed into One Shot MAX Efficiency DH5-T1R cells.

## AUTHOR CONTRIBUTIONS

GK, MD, and JJB initiated the project. JJB conceived *in vivo* recombination strategy. GK and JJB performed *in silico* cloning, designed the study, and wrote the manuscript. GK performed the experiments with the help of MDC, RL, JMA, HP, SS, and AAA. SB performed pathological examination according to clinical criteria. All authors contributed to the final editing.

## ACKNOWLEDGEMENTS

We thank Yuri Voziyanov for the gift of Flp-Cre expression vectors. We thank Barry Stripp for the gift of *mTmG* mice. We thank Liqun Luo for providing the *mTmG* mESCs. We also thank Dawen Cai for mKate2 and dsRed antibodies. We acknowledge support from TJ’s Dream Team HEADing for a Cure, the Samuel Oschin Comprehensive Cancer Institute Cancer Research Forum Award (M.D., J.J.B.), the Board of Governors RMI of Cedars-Sinai (M.D., J.J.B), the Smidt Family Foundation, and the Paul and Vera Guerin Family Foundation. JJB was supported by NIH grant R33 CA202900.

